# Calcium-mediated amyloid co-aggregation of S100A1 and S100A8 proteins

**DOI:** 10.1101/2024.11.26.625466

**Authors:** Viktorija Karalkevičiūtė, Ieva Baronaitė, Aistė Peštenytė, Dominykas Veiveris, Gediminas Usevičius, Mantas Šimėnas, Mantas Žiaunys, Vytautas Smirnovas, Darius Šulskis

## Abstract

The S100 family consists of calcium binding proteins that are largely known for their contribution to the neuroinflammatory processes. They are associated with various cardiac and neurological functions as well as related diseases. A few S100 proteins can form unspecific or amyloid aggregates in neuropathologies and thus play a part in dementia pathogenesis. Among all S100 proteins, S100B and S100A9 aggregation properties are the most investigated, however, there is a lack of studies regarding other S100 members. In particular, S100A1 and S100A8 are also associated with neuropathies, but their interactions or aggregation are poorly understood. Therefore, in this study, we explored whether S100A1 and S100A8 proteins can form heterodimers, interact or co-aggregate.

Our results revealed that S100A1 and S100A8 interactions and amyloid aggregation are driven by calcium ions. We observed that while S100A1 remains mostly stable, S100A8 forms various types of spherical or unspecific aggregates. While they do not form stable heterodimers like calprotectin, their transient interactions facilitate the formation of worm-like amyloid fibrils and the process is regulated by different calcium ion concentrations. At calcium ions saturation, both proteins are stabilized leading to inhibition of aggregation. Overall, by employing a diverse range of techniques from amyloid and protein-specific fluorescence detection to electron-electron double resonance spectroscopy, we elucidated interactions between S100 proteins that might otherwise be overlooked, enhancing our understanding of their aggregation behaviour.

## Introduction

The S100 is a calcium-binding protein family with at least 21 members found in various tissues^1^. They have numerous intracellular and extracellular functions, that range from apoptosis, inflammation, homeostasis to regulating other cells^2^. S100 proteins are found within cells mainly as homodimers, with some key members forming heterodimers for specific functions^3^. The conformation of S100 is controlled by two EF-hand structural motives that bind calcium ions and are essential for S100 functions^4^. Furthermore, the majority of their interactions with their clients are calcium-dependent, allowing them to dynamically participate in calcium signalling pathways^1^. Contributing to that, there is an established connection between calcium ions and S100 oligomerization, as they stabilize dimers, but can also assist in assembling larger oligomers or heterodimers^5^. S100 proteins are located in different parts of the body^2^, however, historically they were first discovered in the brain^6^. Currently, seven members (S100B, S100A1, S100A6, S100A7, S100A8, S100A9, S10012) are known to localize within the brain and are associated with Alzheimer’s disease^7^, regulation of neuroinflammation and neuroactivation^8^ (Fig. 1A).

**Figure 1.**
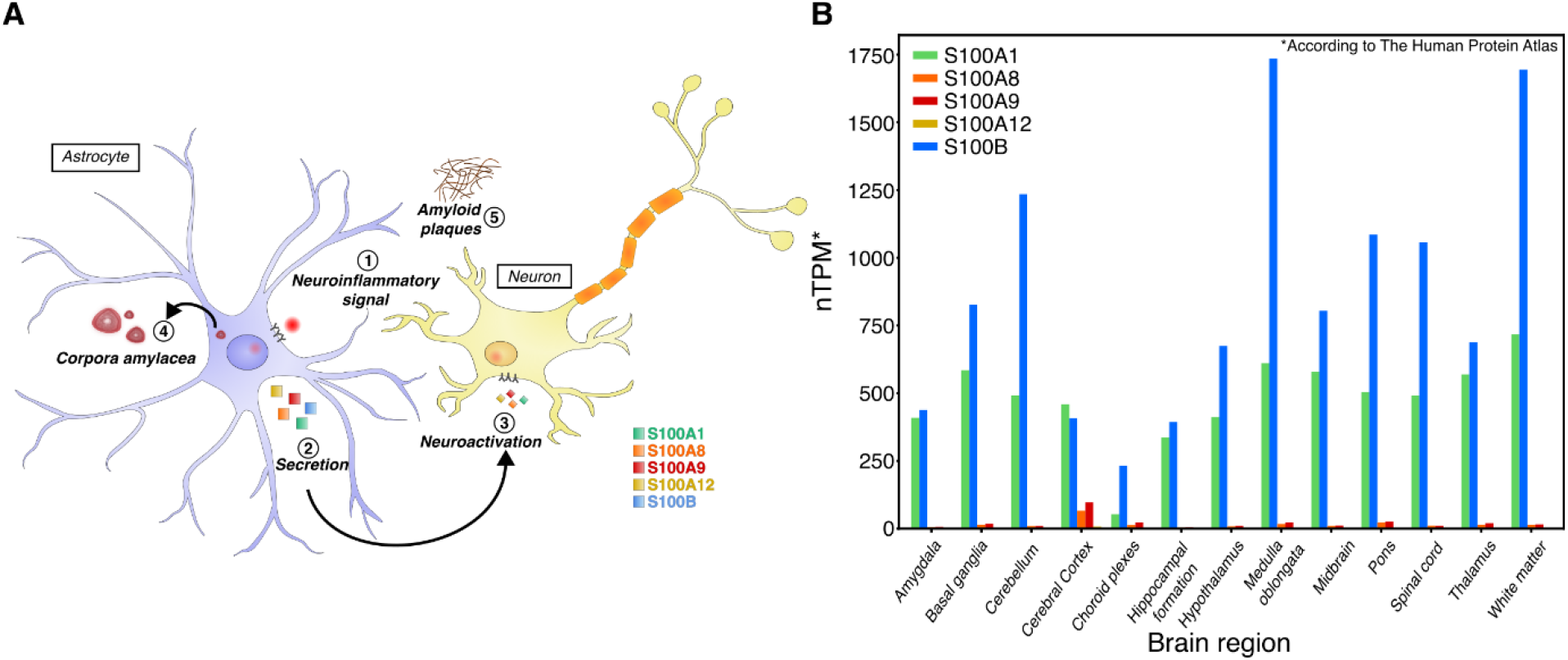
(**A**) During neuroinflammation (1), astrocytes upregulate the expression of S100A1, S100A8, and other S100 proteins^7^. These proteins are secreted via non-canonical pathways^26^ (2) and transfer inflammation signals to other neuronal cells^2^ (3). S100 proteins can accumulate and contribute to the formation of corpora amylacea^11^(4) and amyloid plaques^7^ (5). (**B**) Transcriptional differences of S100 RNA in various brain regions according to The Human Protein Atlas^15^ (nTPM - transcripts per million).

A hallmark feature of various neurodegenerative disorders is insoluble protein aggregates— amyloid fibrils^9^ and a number of S100 proteins form fibrils with distinct worm-like morphologies^10^ Several S100 members can also accumulate in corpora amylacea (CA) - glycoproteinaceous structures that appear in the aging brain and especially during neuropathologies (Fig. 1A). The most abundant S100 proteins in CA are S100A1, S100A8 and S100A9^11^, with S100A8 and S100A9 being shown to form amyloid fibrils or oligomeric complexes that are sensitive to amyloid dyes^12^. Furthermore, S100A8 can form a heterodimer with S100A9 and assemble into calprotectin (CP), which stabilizes both proteins and prevents their aggregation^12^, however, prolonged incubation of CP with zinc/calcium ions induces fibrillation^13^. S100A1 protein is highly prevalent in the heart^14^, however, it is also located in the brain, rivalling transcriptional levels to S100B according to Human Protein Atlas^15^ (Fig. 1B). S100B expression is known to correlate with neurodegenerative disorders^16^ and S100B can form a heterodimer with S100A1^17^, but there is little information about S100A1’s role in neuropathies. S100A1 is expressed in astrocytes^18^ (Fig. 1A), can be found extracellularly^19^ and is involved in neuroinflammation as a calcium ion sensor in Alzheimer’s disease mouse model^20^. Similarly to the S100B and S100A1 protein pair, there is a lot of information about S100A9’s role in neurodegeneration^21,22^, however, there is a substantial lack of studies regarding its interaction partner S100A8. S100A8 is also produced in astrocytes (Fig.1B) and is linked to the enhanced production of beta-amyloid peptide through a positive feedback loop^22^. Recently, we have shown that S100A8 formed various types of aggregates, but not fibrillar structures, unlike several other S100 proteins^10^. In general, only a few other S100A heterodimers are known^17,23–25^, and their possible functions *in vivo* are not clearly understood. In this study, we investigated the interplay between S100A1 and S100A8 proteins and its impact on their stability as well as amyloid aggregation.

Our results revealed that S100A1 does not aggregate and S100A8 forms only heterogeneous aggregates. Upon mixing the two proteins, we observed calcium-concentration-dependent aggregation, resulting in fibril formation. At the intermediate calcium concentration, we observed worm-like fibrils, while higher calcium concentrations inhibited aggregation. Single-molecule microscopy confirmed that both S100A1 and S100A8 co-aggregated to larger aggregate ensembles in the presence of calcium, However, double electron-electron resonance spectroscopy indicated no evident formation of a heterodimer, suggesting that the interactions are transient but substantial enough to affect the aggregation pathways. Overall, we confirmed that the mixture of S100A1 and S100A8 proteins leads to the formation of amyloid fibrils.

## Methods

### Cloning

The SUMO-S100A1 gene was purchased from GENEWIZ (Azenta Life Sciences). The gene was inserted into a pET28a(-) vector via the NdeI and BamHI restriction sites by standard cloning techniques^27^ yielding SUMO-S100A1 construct fused to an amino-terminal His6 tag. Primers for mCherry and eGFP genes were generated by restriction free (RF) cloning tool (https://www.rf-cloning.org)^28^ and RF cloning method^29^ was used to construct mCherry-S100A1 (pVK1) and eGFP-S100A8 (pVK2) plasmids in pET28a(-) backbone vector. Plasmids and primers used in this study can be found in Table S1.

### Protein expression and purification

Plasmid constructs encoding 6xHis-SUMO-S100A1, 6xHis-SUMO-S100A8, 6xHis-mCherry-S100A1 and 6xHis-eGFP-S100A8 were transformed into One Shot™ BL21 Star™ (DE3) *E. coli* (Thermo Scientific) cells by heat shock (42°C 45 s). Transformed cells were grown in 100 mL of LB medium containing kanamycin (50 μg/mL) at 37°C, 220 rpm, for 16 hours. The culture was transferred to 200 mL of LB medium with kanamycin (50 μg/mL) and grown at 37°C, 220 rpm until optical density at 600 nm reached 0.6–0.8. Protein expression was induced by adding 200 μM IPTG and the culture was incubated at 25°C, 220 rpm, for 18 hours. Cells were harvested by centrifugation (6000 g, 20 min, 4 °C). The biomass was resuspended in buffer (25 mM HEPES, 1.0 M NaCl, 10 mM imidazole, pH 8.0), which was followed by the addition of lysozyme and 1 mM phenylmethylsulfonyl fluoride (PMSF). Cells were lysed by sonication (Sonopuls, VS70T probe; Bandelin) for 30 min at 40% amplitude, with a 15-second on/30-second off cycle. The lysate was centrifuged (18,000 rpm, 30 min., 4 °C) and the supernatant was filtered through a 0.45 μm pore size filter.

Protein purification via immobilized metal ion affinity chromatography (IMAC) was performed using a gravity column packed with Ni^2+^ Sepharose 6 Fast Flow resin (Cytiva). The column was washed with 50 mM HEPES, 1.0 M NaCl, 10 m imidazole (pH 8.0) buffer, followed by elution using the same buffer solution that contained imidazole concentrations of 0.05 M and 0.5 M imidazole. For all samples the last fraction was collected.

6xHis-SUMO-S100A1 and 6xHis-SUMO-S100A8 were dialysed (8000 MWCO, Biodesign™ D106) in 10 mM Tris buffer (pH 8.0) for 1 hour. Sentrin-specific protease 1 (SENP1) was added to cleave the 6xHis-SUMO tag, and the samples were dialyzed for 18 hours in fresh 10 mM Tris buffer (pH 8.0). Then the samples were centrifuged (3160 g, 30 min, 4°C) and filtered through a 0.45 μm pore size filter. Purification via IMAC was repeated to collect the flow-through. All fractions were checked by SDS-PAGE. Before size exclusion chromatography, 10 mM of EDTA and dithiothreitol (DTT) was added to all the samples (S100A1, S100A8, 6xHis-eGFP-S100A8, 6xHis-mCherry-S100A1). The samples were then concentrated (10 kDa MWCO, Merck) and filtered through a 0.22 μm pore size filter. Size exclusion chromatography was performed with columns packed with Superdex™ 75 sorbent (Cytiva) for S100A1 and S100A8 or Superdex™ 200 sorbent (Cytiva) for 6xHis-eGFP-S100A8 and 6xHis-mCherry-S100A1 proteins, calibrated with <u>50 mM HEPES (pH 7.4) buffer</u>. The collected fractions were checked by SDS-PAGE and concentrated (10 kDa MWCO, Merck). The protein samples were stored at -80 °C.

### Differential Scanning Fluorimetry (DSF)

Samples for the protein stability assay contained 100 μM of either S100A1, S100A8, or both S100A1/A8 prepared in 1 mM TCEP, 50 mM HEPES (pH 7.4) buffer with increasing CaCl_2_ concentrations (0, 50, 100, 200, 400, 800, 1600 μM) supplemented with 100 μM 8-anilinonaphthalene-1-sulfonic acid (ANS). The ANS concentration was determined its extinction coefficient (ε_351nm_ = 5500 M^-1^cm^-1^). For the sample without CaCl_2_, 1 mM EDTA was added. Additionally, S100A1 samples were prepared containing 0.5, 1.0 M guanidinium chloride, without CaCl_2_. Protein unfolding was monitored with a Rotor-Gene Q instrument (QIAGEN) using the blue channel (excitation, 365 ± 20 nm; detection, 460 ± 20 nm). The unfolding process was initiated by ramping the temperature from 25°C to 99°C at 1°C/min increment. Data analysis was performed using MoltenProt software^30^.

### Nano Differential Scanning Fluorimetry (nanoDSF)

S100A1 protein was desalted using a desalting column (BioWorks) with a 50 mM TRIS buffer (pH 7.5). Prometheus NT.48 Series nanoDSF grade standard capillaries (NanoTemper Technologies) were filled with a sample containing 100 μM S100A1. Protein unfolding was monitored using a Prometheus NT.48 instrument (NanoTemper Technologies) by measuring absorbance at 330 and 350 nm (with 20% excitation). The temperature was increased from 20°C to 99°C at a rate of 1°C/min. Data analysis was performed using MoltenProt software^30^.

### Thioflavin T (ThT) fluorescence assay

The amyloid-specific Thioflavin-T (ThT) dye was used to monitor aggregation kinetics. Samples for the ThT fluorescence assay contained 1 mM TCEP, 50 μM ThT, and 100 μM of either S100A1, S100A8 or S100A1/A8 in 50 mM HEPES buffer (pH 7.4). Each sample was prepared with increasing concentrations of CaCl_2_ (0, 50, 100, 200, 400, 800, 1600 μM). For the sample containing no CaCl_2_, 1 mM EDTA was added. 100 μL of each sample was placed into three separate wells of a 96 well non-binding plate. Aggregation kinetics were measured every 5 min at 42°C using a CLARIOstar Plus microplate reader. ThT dye was excited at 440 nm and the emission signal was recorded at 480 nm.

### Fourier-transform infrared (FTIR) spectroscopy

After 65 h aggregation at 42°C, samples were removed from the aggregation reaction kinetics plate and were used for the preparation of FTIR measurements (280 μL of each sample). The aggregated samples of S100A8 were centrifuged at 16,900 g for 30 min, after which the supernatant was removed and replaced with 300 μL D2O supplemented with 400 mM NaCl (addition of NaCl may improve fibril sedimentation^31^. The centrifugation and resuspension procedures were repeated four times. After the final step, the aggregate pellet was resuspended into 50 μL D2O containing 500 mM NaCl.

FTIR spectra were acquired as described previously^32^ using a Invenio S FTIR spectrometer (Bruker), equipped with a liquid-nitrogen-cooled mercury-cadmium-telluride detector, at room temperature and constant dry-air purging. For every sample, 256 interferograms of 2 cm^-1^ resolution were recorded and averaged. D2O containing 400 mM NaCl and water vapor spectra were subtracted from each sample spectrum, followed by baseline correction and normalization to the same 1,595-1,700 cm^-1^ wavenumber range. All data processing was done using GRAMS software.

### Far-UV circular dichroism (CD) spectroscopy

Samples for CD spectra measurements contained 100 μM of either S100A1, S100A8, or S100A1/A8 in a 1 mM TCEP, 50 mM HEPES buffer (pH 7.4). Each sample was prepared with increasing concentrations of CaCl_2_ (0, 50, 100, 200, 400, 800, 1600 μM). For the sample without CaCl_2_, 1 mM EDTA was added. Aggregation of samples was conducted in 1.5 ml microcentrifuge tubes (Eppendorf) at 42°C for 70 hours. The samples were then placed in a 0.5 mm quartz cuvette, and CD spectra were measured using a J-815 spectropolarimeter (Jasco). For each sample, spectra between 190 and 260 nm were recorded at 0.2 nm intervals. The data was smoothed using Gaussian smoothing (SD=10). All data processing was done using Quasar^33^.

### Atomic force microscopy (AFM)

The freshly cleaved mica was positively charged by applying 50 μL of 0.5% (3-Aminopropyl)triethoxysilane (APTES) on the surface and allowing it to functionalize for 5 min. Subsequently, mica was washed with dH_2_O, dried with airflow and the procedure was repeated with 50 μL of the protein sample. Imaging was performed using a Dimension Icon microscope (Bruker) operating in tapping-in-air mode with aluminium-coated silicon tips (RTESPA-300, Bruker). The images were processed using Gwyddion 2.66 software^34^. The cross-sectional height of aggregates was determined from extracted line profiles, which were fitted using the Gaussian function. The examples of cross-sectional height profiles are presented in supplementary data (Fig. S1).

### Fluorescence Microscopy

The samples for fluorescence microscopy were prepared as in ThT fluorescence assay with the exclusion of ThT dye and the addition of 1 μM of either or both mCherry-S100A1 and eGFP-S100A8 proteins, resulting in 99 μM non-tagged, 1 μM tagged proteins respectively.

15 μL aliquots of each sample were pipetted onto 1 mm glass slides (Fisher Scientific, cat. No. 11572203), covered with 0.18 mm coverslips (Fisher Scientific, cat. No. 17244914) and imaged as described previously^35^ using an Olympus IX83 microscope with a 40x objective (EVIDENT, NA 0.6, LUCPLFLN40X) and fluorescence filter cubes (475-495 nm excitation and 510-550 nm emission wavelengths for eGFP-S100A8, 540-550 nm excitation and 575-625 nm emission wavelengths for mCherry-S100A1). Images were captured using an ORCA-Fusion Digital CMOS camera (Hamamatsu, model C14440-20UP). Data analysis was done using Fiji software^36^. Not-cropped fluorescence images are presented in Fig. S2-4.

### Transmission electron microscopy (TEM)

On glow-discharged 300-mesh formvar/carbon supported copper grids (Agar Scientific), 5 μL of each sample prepared for fluorescence microscopy was applied, incubated for 1 min, and then the grid was dried with filter paper. The same procedure was repeated with 5 μL of 2% (w/v) uranyl acetate, followed by two washes with 5 μL of dH_2_O. TEM images were acquired using a Talos 120C (Thermo Fisher) microscope operating at 120 kV, equipped with a 4k × 4k Ceta CMOS camera. Images were processed using Fiji^36^.

### Chaperone activity assay

Samples containing 10 μM of S100A1, S100A8, and S100A1/A8 with 0.2 mg/ml lysozyme, were prepared in a buffer solution of 20 mM DTT and 50 mM HEPES (pH 7.4). Three samples were prepared with increasing concentrations of CaCl_2_ (0, 20, 200 μM). For the sample that did not contain CaCl_2_, 2 mM EDTA was added. 100 μL of each sample was placed into four separate wells of a 96-well non-binding plate. Chaperone activity was measured spectrophotometrically by monitoring absorbance at 360 nm using a CLARIOstar Plus microplate reader at 37°C.

### Alpha-fold prediction

Protein structures were predicted using AlphaFold3^37^. The amino sequences of S100A8 and S100A1 with or without 4 atoms of calcium ions were used as input and run with default settings. S100A8/S100A1 confidence scores were ipTM = 0.8 and pTM = 0.81 in the presence of calcium ions. Confidence levels and scores are depicted in Fig. S5.

### Electron paramagnetic resonance (EPR)

For electron paramagnetic resonance (EPR) double electron-electron resonance (DEER) measurements proteins were labelled as described previously^38^. Briefly, S100A1 or S100A9 were incubated with 10 mM DTT for 1 hour and desalted using a BabyBio desalting column (Bio-Works) with PBS. Next, proteins were incubated with 10 times excess of nitroxide spin label MTSSL (Sigma-Aldrich) reagent overnight at 8°C with gentle shaking. Proteins were desalted again to 50 mM HEPES 10 % Glycerol (pH 7.4) buffer in the morning. The final concentrations of labelled S100A1 and S100A9 proteins were 50 μM and 100 μM respectively. For heterodimer measurements, proteins were mixed equimolarly with their partners.

DEER spectroscopy measurements were performed at the X-band (9.5 GHz) microwave frequency using a Bruker ELEXSYS E580 EPR spectrometer. The sample was cooled to 50 K using liquid helium, in a helium flow cryostat. A 4 mm diameter sample tube containing approximately 20 μL of sample was flash-frozen in liquid nitrogen and subsequently inserted into a Bruker ER4118X-MD5 microwave resonator. For improved sensitivity, a cryoprobe equipped with a cryogenic low-noise microwave amplifier was employed^39^.

An echo-detected field-sweep EPR spectrum was recorded using a Hahn-echo pulse sequence. The pulse sequence for the four-pulse DEER experiment was:

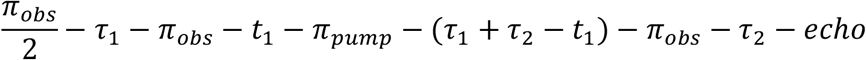

Measurements were acquired with an interpulse delay τ_1_ of 400 ns for S100A1 and 300 ns for S100A9 samples. The dead time delay t_1_ of 80 ns was used for all samples, while the interpulse delay τ_2_ and the shot repetition time were optimized for each sample. The microwave power was optimized to obtain a π/2-pulse length of 16 ns and a π-pulse length of 32 ns for both pump and observer pulses. All traces were collected using eight-step nuclear modulation averaging, with an averaging time step of 56 ns, a 20 ns time-domain for the primary data, and 60 MHz frequency difference between the observer and pump pulses. The data analysis was performed using DeerLab^40^ (release 1.1.2), a Python-based package for DEER spectroscopy data analysis. The experimental signal was modeled using the general kernel, assuming a homogeneous spin distribution in a 3D medium for the background. The distance distribution was fitted using a Tikhonov regularization, with the regularization penalty weight selected using the Akaike information criterion and a non-negativity constraint. The asymptotic method was used to determine the 95% confidence intervals. The DeerLab results were also compared to other software tools, including DeerAnalysis 2022^41^ and DeerNet^42^, all of which produced highly consistent outcomes.

## Results and Discussion

### S100A1 alters aggregation of S100A8

The aggregation of S100A1 and S100A8 was followed by the amyloid-specific fluorescent ThT dye^43^. Aggregation was not observed for S100A1 with or without calcium ions (Fig. 2A), whereas S100A8 aggregated in a two-phase manner and calcium extended the duration of the first phase (Fig. S6), consistent with previous findings^12^. When S100A1 and S100A8 were mixed in equimolar concentrations, the aggregation kinetics changed. At CaCl_2_ concentrations up to 50 μM, proteins aggregated in a bi-phasic manner, but at higher calcium ion concentrations, the aggregation transitioned to an exponential growth phase, resembling the kinetics observed for S100A9^44^. Comparing the maximum ThT fluorescence levels at the end of the aggregation process (Fig. 2B), S100A8 aggregation fluorescence exponentially decreased with increasing calcium ion concentration, but the intensities of the S100A1/S100A8 samples were higher, presumably due to calcium ions binding competition between S100A1 and S100A8 proteins. At the highest 1600 μM CaCl_2_ concentration, aggregation was strongly inhibited, hinting that both calcium-binding EF-Hands of S100A1 and S100A8 were saturated with calcium ions, leading to stabilized native structures.

**Figure 2.**
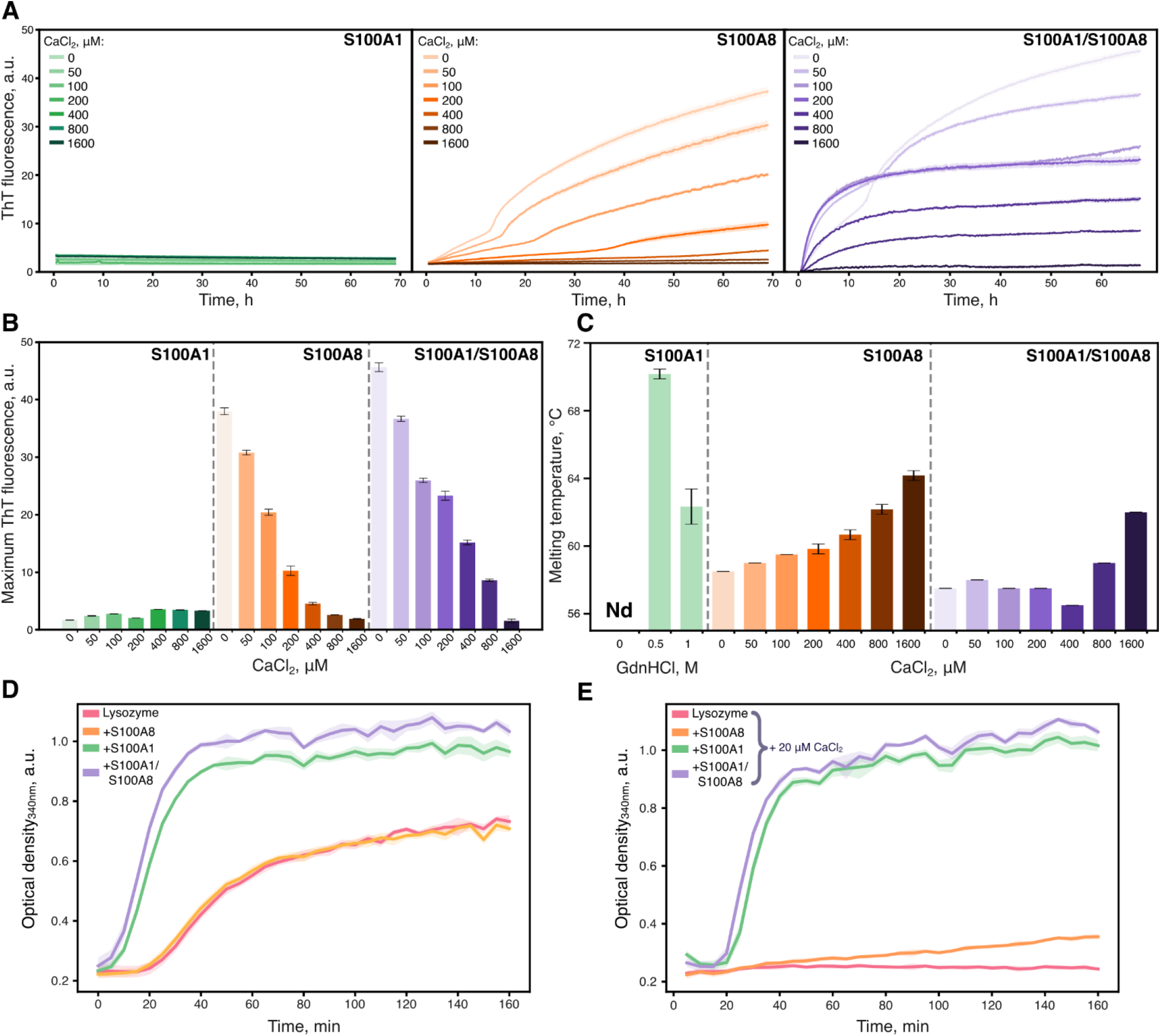
Aggregation and stability propensities of S100A1 and S100A8 proteins (**A**) Aggregation kinetics of S100A1, S100A8 and S100A1/S100A8 in the presence or the absence of calcium ions, followed by ThT fluorescence. (**B**) Maximum ThT fluorescence values of aggregation kinetics. (**C**) Melting temperatures of S100A1, S100A8 and S100A1/S100A8 proteins. Chaperone assay of S100A1 (10 μM), S100A8 (10 μM) and S100A1/S100A8 (10 μM of each) against DTT-induced aggregation of lysozyme (0.2 mg/ml) in the absence (**D**) and the presence of CaCl_2_ (20 μM) (**E**).

Concurrently, we investigated the protein melting temperature using differential scanning fluorimetry by measuring the fluorescence of the ANS dye (Fig. S7), which binds to hydrophobic pockets upon protein unfolding or aggregation^41^. We did not observe the unfolding of S100A1 under initial conditions, only upon the addition of a denaturing reagent (guanidine hydrochloride), noticeable melting temperatures were obtained (Fig. 2C). This contrasts with previously reported S100A1 melting temperatures, which ranged from 70 to 75 ºC when measured in Tris buffer without guanidine hydrochloride using nanoDSF^45^. Therefore, we repeated the measurement under those conditions (Fig. S8) and observed S100A1 melting at a slightly higher temperature of 85 ºC, which might be due to different protein preparations. Since nanoDSF measures protein changes via aromatic residue shifts^46^, there is a possibility that it detects intrinsic conformational changes or partial unfolding, which DSF does not detect. However, we did observe denaturation of S100A8 using DSF and calcium ions further stabilised S100A8 as reported previously^12,47^. In the mixture of both proteins, melting temperatures were slightly lower and increased at higher calcium ion concentrations (>400 μM), suggesting potential competition for calcium binding or transient interactions between the proteins.

We also investigated whether S100A1 exhibits any chaperone activity that could influence aggregation since it is known to form a multi-chaperone complex with Hsp70/90^48^, and other S100 members also have been reported to have chaperone-like functions^49,50^. In the lysozyme chaperone activity assay (Fig. 2D), S100A1 or a mixture of both proteins accelerated the aggregation of lysozyme, consistent with previous reports for S100A6^51^, whereas S100A8 had no effect. Notably, the presence of calcium ions inhibited the aggregation of lysozyme (Fig. 2E), but in the mixture with S100A1 or S100A1/S100A8, aggregation was observed, due to calcium ions being salvaged by S100A proteins. In higher calcium concentrations (Fig. S9), lysozyme aggregation was inhibited under all conditions, preventing further analysis of chaperone-like activity. These findings suggest that unlike expected chaperone activity, S100A1 interactions may promote co-aggregation or modify aggregation pathways when in complex with other proteins.

### S100A1/S100A8 amyloid fibrillation is dependent on calcium concentration

We investigated the aggregates formed by S100A1 and S100A8 using three different microscopy techniques. To begin with, atomic force microscopy revealed distinct morphologies of assemblies. S100A1 formed a small number of amorphous aggregates, whereas S100A8 aggregated into spherical oligomers/clusters (Fig. 3A). Upon the addition of calcium ions, the number of precipitates in the S100A8 sample decreased, correlating with the lower fluorescence of ThT in the aggregation kinetic data (Fig. 2A) and previously published results^12^. Although in the absence of calcium ions S100A1/S100A8 resembled S100A8 aggregates, worm-like fibrils started to form upon the addition of calcium ions (Fig. 3B). The size of the aggregates steadily increased from 2.14 nm to 2.86 nm with calcium ion concentrations up to 400 μM (Fig. 3C). At higher than 400 μM CaCl_2_, fewer aggregates were observed, likely related to overall protein stabilization. Worm-like fibrils indicate that S100 proteins potentially only partially unfold, leading to curly fibrils, similar to lysozyme fibrils^52^. To support observations by AFM, additional images with transmission electron microscope (Fig. 3D) were acquired using higher concentrations of proteins. Samples of S100A1 and S100A8 either contained no aggregates or formed amorphous assemblies, respectively (Fig. S10A). At 200 μM CaCl_2_ worm-like fibrils were observed, in correspondence with AFM results. Surprisingly, at the highest CaCl_2_ concentration (1600 μM), fibrils were also detected, although they seemed more likely to cluster.

**Figure 3.**
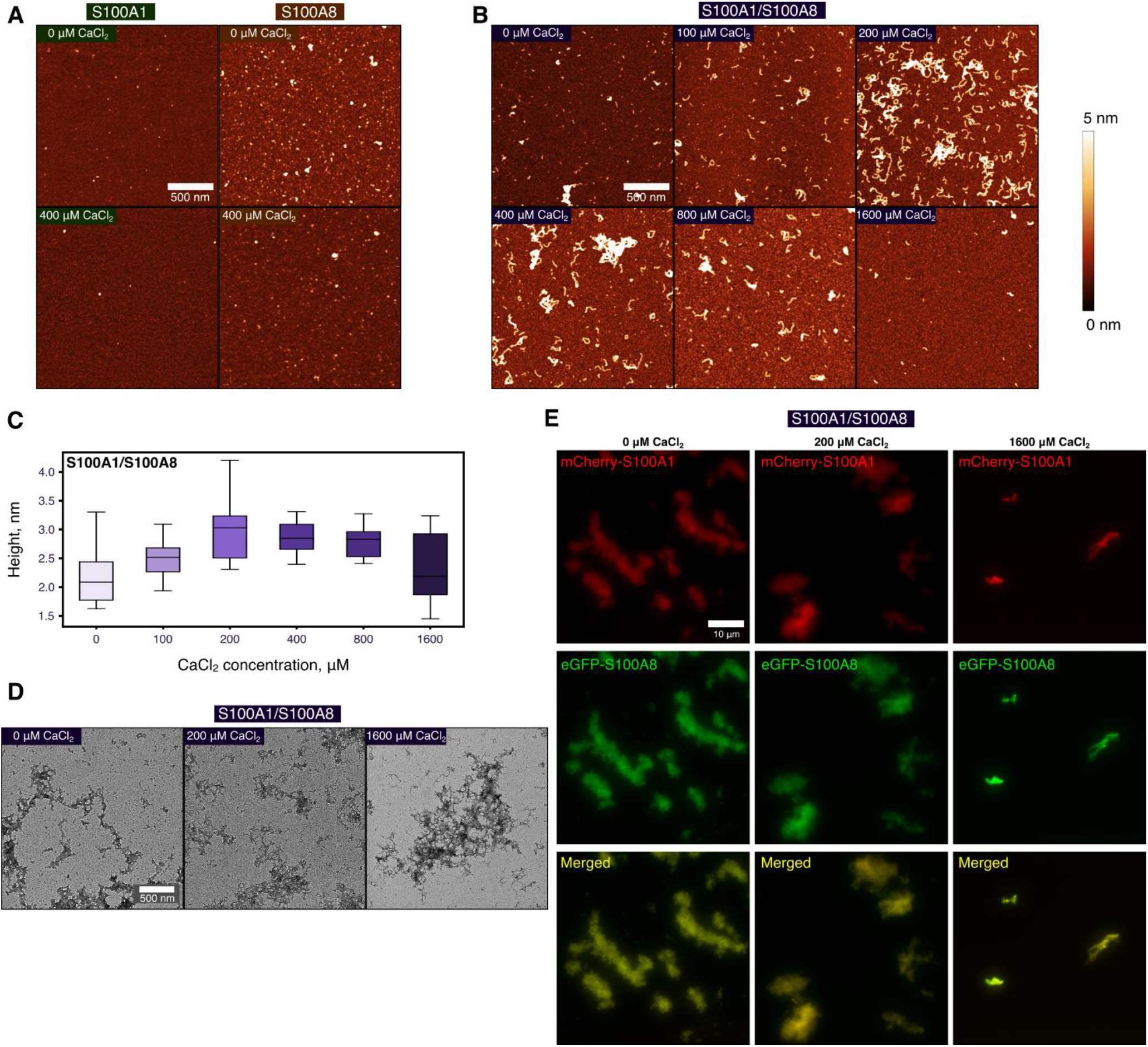
Morphology and co-localization of S100A1 and S100A8 aggregate. AFM images of (**A**) S100A1, S100A8 and (**B**) S100A1/S100A8 after 65h of aggregation (scale bar 500 nm). (**C**) The height distribution of S100A1/S100A8 samples with box plots indicating the median, interquartile range (IQR) and whiskers 1.5x range of IQR from box. (**D**) The transmission electron microscopy (scale bar 500 nm) and (**E**) fluorescence microscopy (scale bar 10 μm) images of S100A1/S100A8 aggregated mixtures at different calcium ion concentrations. Additional fluorescence images are presented in Fig. S2-4.

Finally, fluorescence microscopy was employed to investigate whether S100A1 and S100A8 co-localize (Fig. 3E, S10B). At all range of calcium concentrations, we observed co-localization of both proteins in large plaques. The largest difference was at 1600 μM CaCl_2_, where only much smaller and fewer clumps were seen, which is expected due to aggregation inhibition. Altogether, in the mixture of both proteins, worm-like fibrils are formed and both proteins can co-localize, even at the highest calcium concentration.

### Structural properties of S100A1/S100A8 complex

After the imaging, we conducted structural investigations of S100A1 and S100A8 proteins. Firstly, we examined their structure post-aggregation using circular dichroism. S100A1 exhibited globular conformation in all conditions with two minimums at 209 and 222 nm (Fig. 4A), indicating α-helical structure^53^. S100A8, in the absence of calcium ions, displayed aggregation into β-sheets. This was further confirmed by FTIR spectroscopy (Fig. S11), showing an amide I band at ∼1620 cm^-1 54^, in correspondence with previously reported S100A8 and S100A9 aggregates^12,44^ Addition of calcium ions, steadily stabilized S100A8 into α-helical fold. Although we observed reduced ellipticity for the mixture of both proteins in lower calcium ion concentration ranges, the spectra consistently resembled α-helical structure throughout all tested conditions.

**Figure 4.**
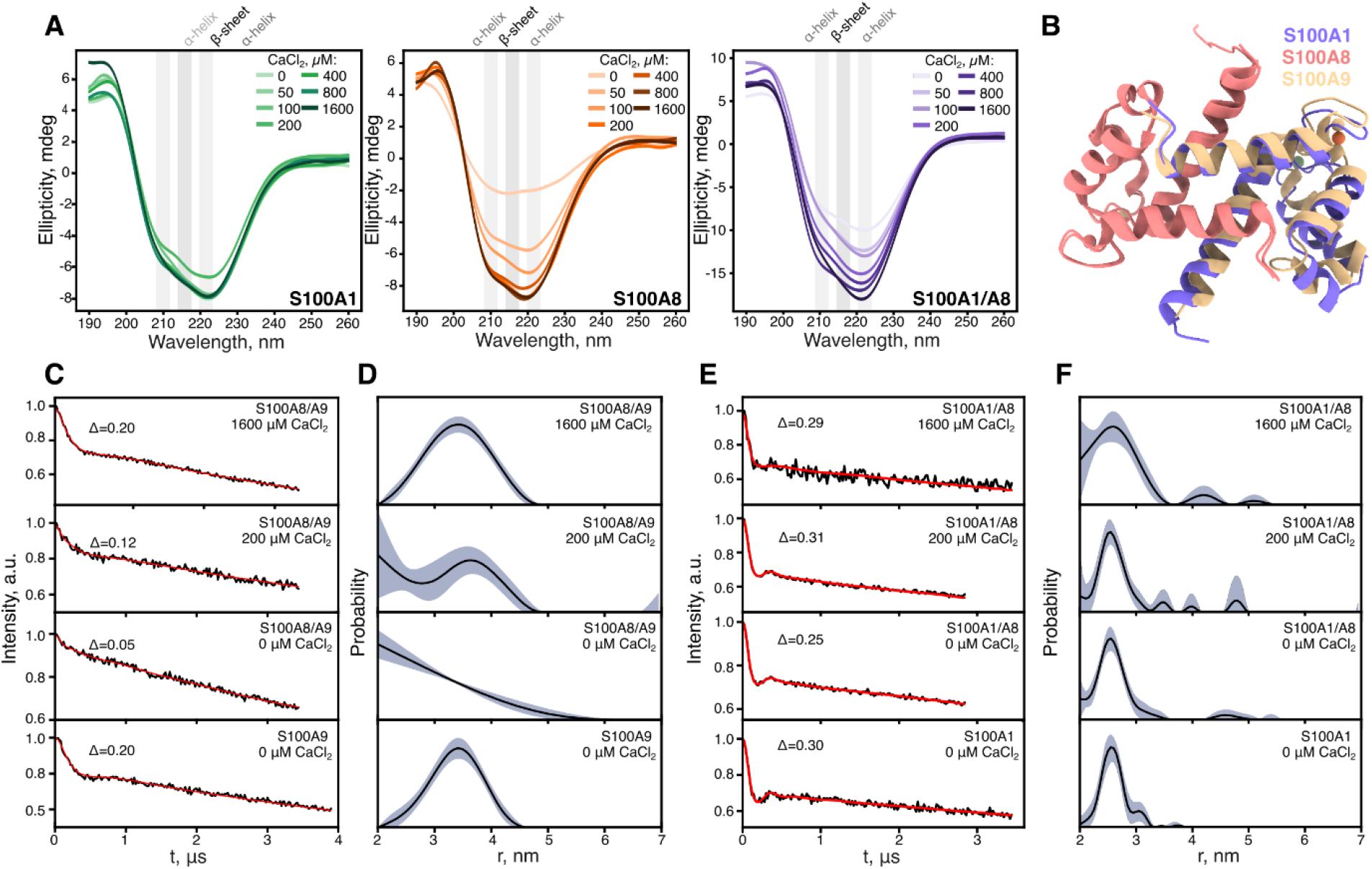
Structure and cross-interaction of S100A1 and S100A8 proteins. (**A**) CD spectra of S100A1, S100A8 and mixture of both proteins after 70 h of aggregation at 42 °C. (**B**) Predicted AlphaFold model of S100A1 and S100A8 heterodimer compared to S100A8/S100A9 heterodimer (PDB id: 1XK4). Time-domain DEER signals and regularized distance distributions of S100A8/S100A9 (**C, D**) and S100A1/S100A8 (**E, F**) samples at different calcium concentrations.

To determine potential heterodimer formation between S100A1 and S100A8, we predicted their structure using AlphaFold 3 webserver^37^ and compared them to the known S100A8/S100A9 heterodimer (Fig. 4B). In both heterodimers, S100A8 maintained an identical conformation, while S100A1 adopted a structure similar to S100A9, with minor alterations in the loops. On the whole, the hypothetical heterodimer of S100A1/A8 resembled S100A8/S100A9.

To further investigate the potential formation of S100A1/S100A8 heterodimer and compare it to characterized S100A8/A100A9, we used DEER spectroscopy. With DEER, we measured distances of spin-labelled cysteines in S100A9 and S100A1 proteins, which were mixed with non-labelled S100A8. The cysteine of S100A9 is located at the N-terminus and S100A1 at the C-terminus, thus allowing accurate measurement of protein diameters (Fig. S12). First, we observed 3.45 nm with FWHM of 0.5 nm distance for S100A9 homodimer and no signal for S100A8/S100A9 complex (Fig. 4C, D), which corresponds to the successful formation of the heterodimer. The signal was recovered with the addition of calcium, indicating dissociation of heterodimers. It is known that S100A8/S100A9 forms tetramers in the presence of calcium ions, however, it requires refolding of both proteins at the same time as mixing them is not sufficient^55^, a process we performed in this study. Overall, by employing DEER spectroscopy, we were able to detect S100A8/S100A9 heterodimer, thus we followed up experiments with S100A1 and S100A8. In all conditions (Fig. 4E, F), we observed distances of 2.55 nm, which is close to S100A1 homodimer diameter (Fig. S12). Altogether, even though AlphaFold3 predicted S100A1/S100A8 heterodimer, S100A1 and S100A8 do not form dimers or their population is too low to be detected by DEER spectroscopy, therefore the likely interaction during co-aggregation is between different homodimers or in larger oligomeric states.

## Conclusions

The S100 family consists of small proteins with a basic structure, yet they exhibit a plethora of functions^2^ and play a major role in activating neuroinflammation, as well as the progression of diseases^8^. In correlation with their large expression in the brain, they have been identified to co-aggregate or interact with proteins associated with neurodegenerative disorder related proteins, such as amyloid-beta^56^, alpha-synuclein^57^ or tau^49^. Despite this, relatively little is known regarding the interactions among different family members, although they are known to oligomerize and form heterodimers^47,58,59^. Furthermore, there is even less understanding of how transient or weak interactions can determine protein structure or aggregation pathways.

In this study, we have shown that S100A1 affects S100A8 aggregation. Unlike S100A8 or S100A9, S100A1 is highly stable and aggregates minimally, however, it can interact with S100A8, which leads to co-aggregation and the formation of worm-like fibrils. Calcium ions further mediate these interactions from promoting to inhibiting aggregation at the highest concentration. Inside the cell, calcium levels are around 100 nM^60^, while extracellular concentrations are approximately 1 mM^61^, therefore, co-aggregation might occur when S100 proteins are released into the extracellular space. On the other hand, during calcium influx, the calcium concentration can increase up to 1 μM and higher ranges^62^, enabling interactions between S100 proteins. These interactions are difficult to observe, but we have identified that they might be critical to protein stability and structural changes.

Another conclusion of the results is that, similarly to moonlight proteins^63^, which catalyse biochemical processes through weak protein or surface interactions, S100A1 allows the formation of S100A8 worm-like fibrils. While S100A1 is known to have co-chaperone activity^48^ which promotes interactions with aggregating proteins such as S100A8, it is insufficient to fully suppress aggregation. Instead, it inhibits amorphous aggregation of S100A8 while promoting amyloid formation. S100A1 does not appear to protect against aggregation, leading to anti-chaperone activity and co-localization. Our study provides a new perspective on S100A1 and S100A8 protein interaction and expands our understanding of them, which have yet to be sufficiently investigated compared to their other protein partners.

## Acknowledgements

We thank Aurimas Kopūstas and Dr. Marijonas Tutkus (Vilnius University) for suggestions and support related to fluorescence microscopy methodology.

## Author Contribution

**Darius Šulskis**: Conceptualization, Formal Analysis, Investigation, Methodology, Writing - Original Draft, Writing -Review & Editing, Visualization. **Viktorija Karalkevičiūtė -** Formal Analysis, Investigation, Methodology, Writing - Original Draft, Writing -Review & Editing, Visualization. **Ieva Baronaitė**: Formal Analysis, Investigation, Methodology, Writing - Original Draft, Writing -Review & Editing, Visualization. **Aistė Peštenytė -** Formal Analysis, Methodology, Writing -Review & Editing, Visualization. **Gediminas Usevičius** -**-** Formal Analysis, Methodology, Writing -Review & Editing, Visualization. **Mantas Šimėnas -** Formal Analysis, Methodology, Writing -Review & Editing, Visualization. **Dominykas Veiveris -** Formal Analysis, Methodology, Writing -Review & Editing. **Mantas Žiaunys**: Formal Analysis, Investigation, Methodology, Writing -Review & Editing. **Vytautas Smirnovas**: Supervision, Writing -Review & Editing.

## Data and materials availability

All data needed to evaluate the conclusions in the paper are present in the paper and/or Supplementary Materials. The raw data used in this paper have been tabulated and are available on Mendeley Data: 10.17632/rr75p7h9f7.1. All other relevant data are available from the corresponding author upon reasonable request.

