## Supplementary material for "Calcium-mediated amyloid co-aggregation of S100A1 and S100A8 proteins"

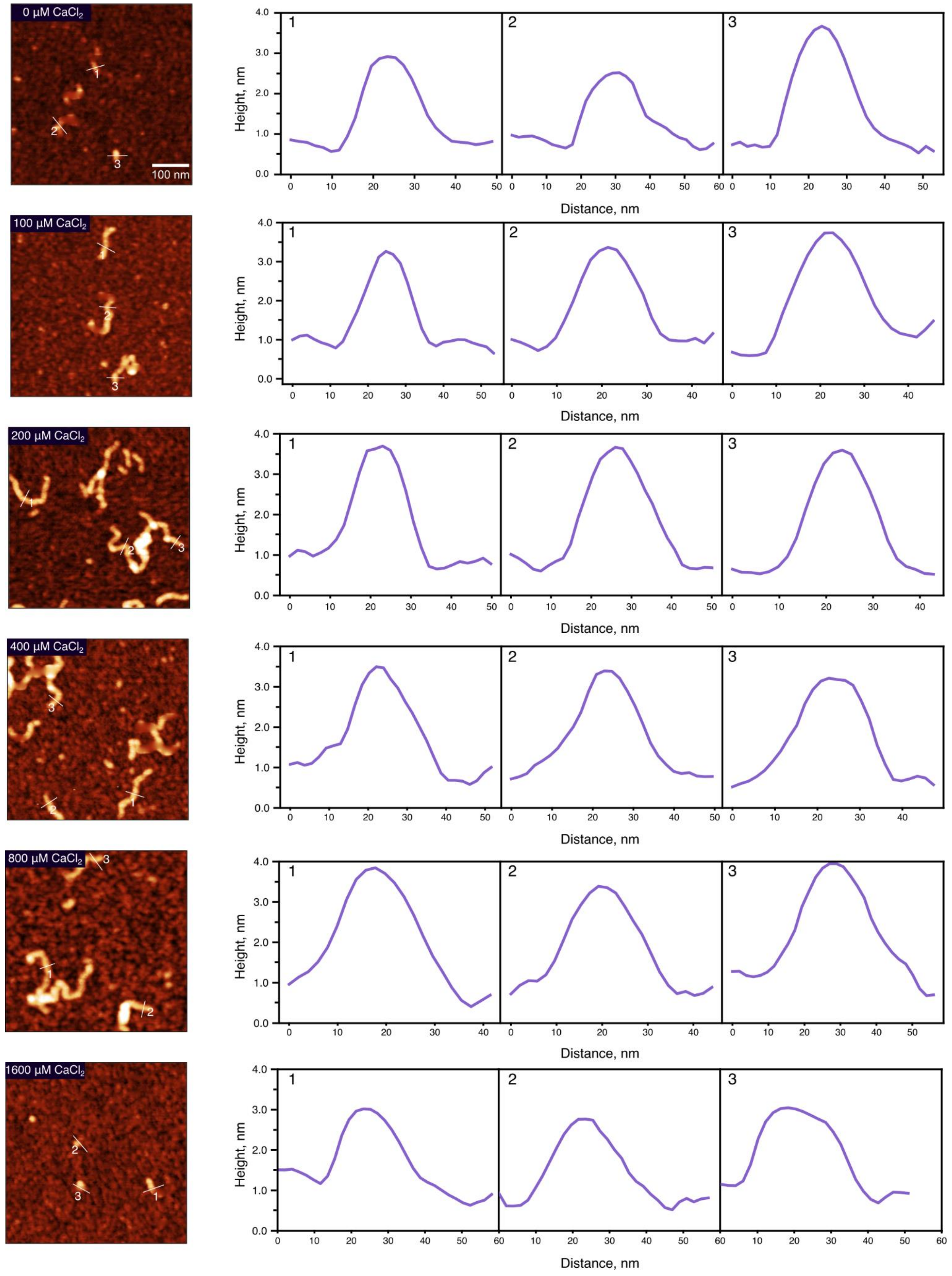


**Figure S1.** The representative height profiles of S100A1/A8 particles from AFM i      mages (scale bar 100 nm).

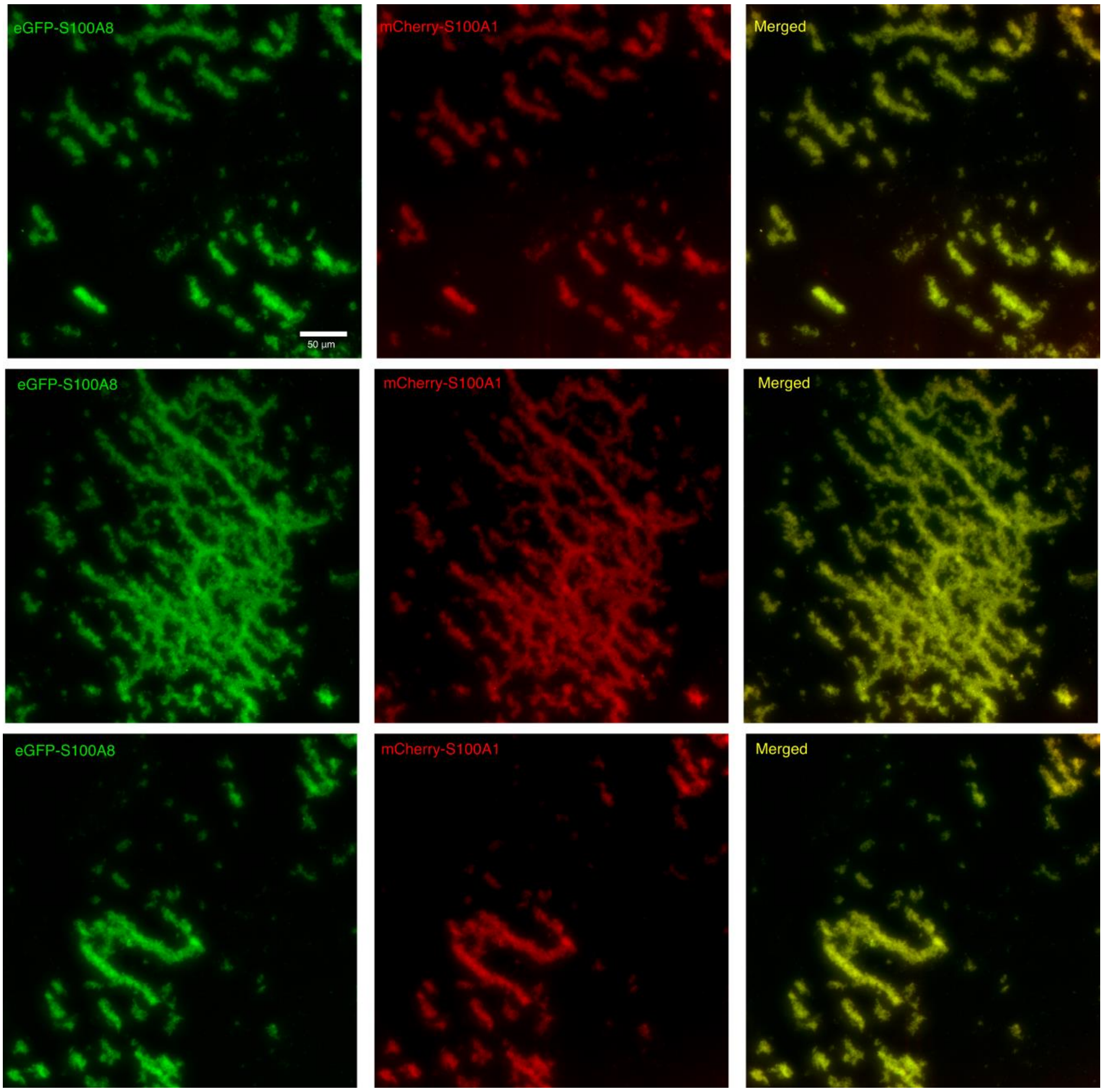


**Figure S2.** Fluorescence microscopy images of S100A1/S100A8 sample after aggregation without $CaCl_2$. Scale bar is 50 μm.

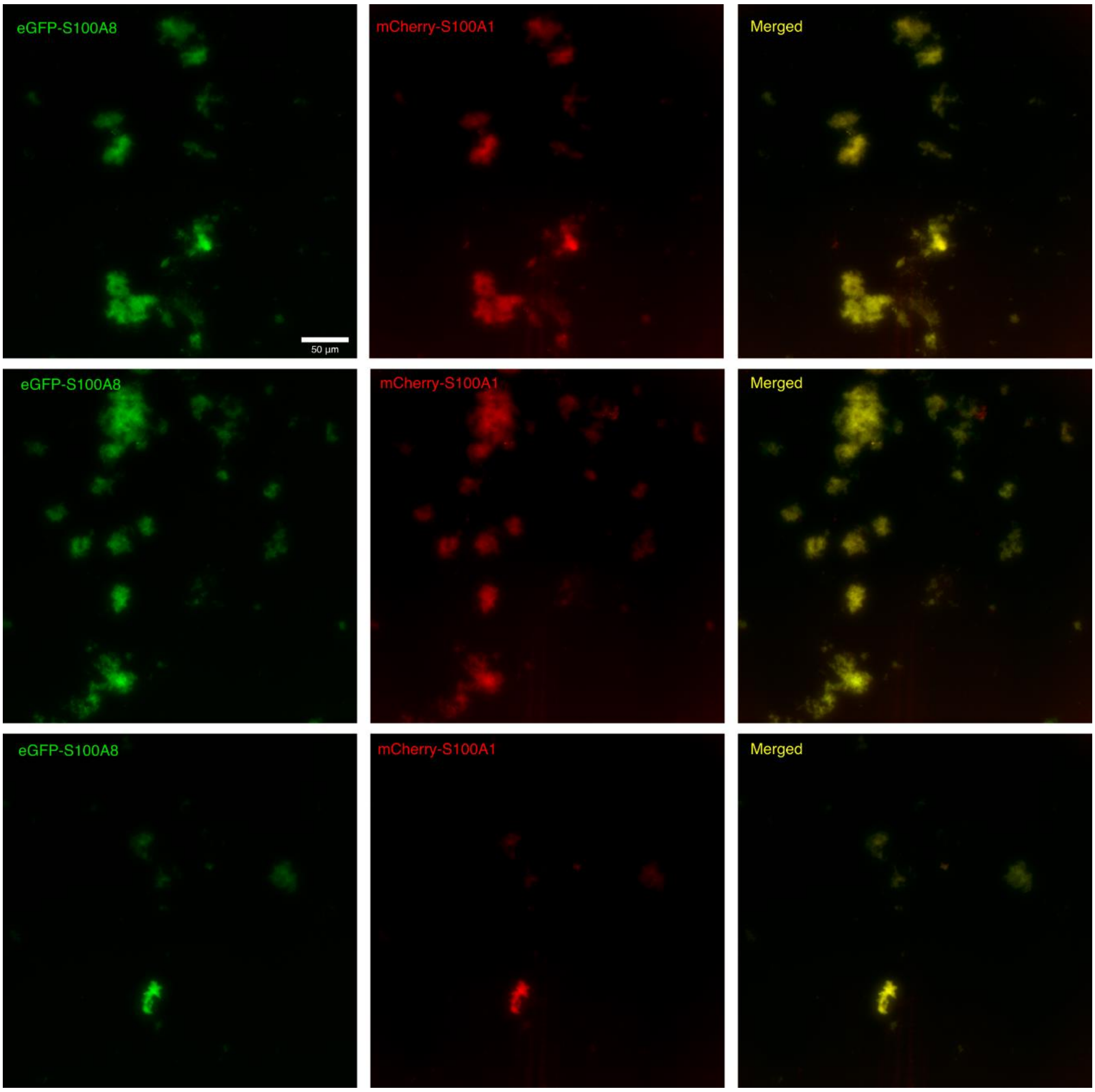


**Figure S3.** Fluorescence microscopy images of S100A1/S100A8 sample after aggregation in the presence of 200 µM $CaCl_2$. Scale bar is 50 µm.

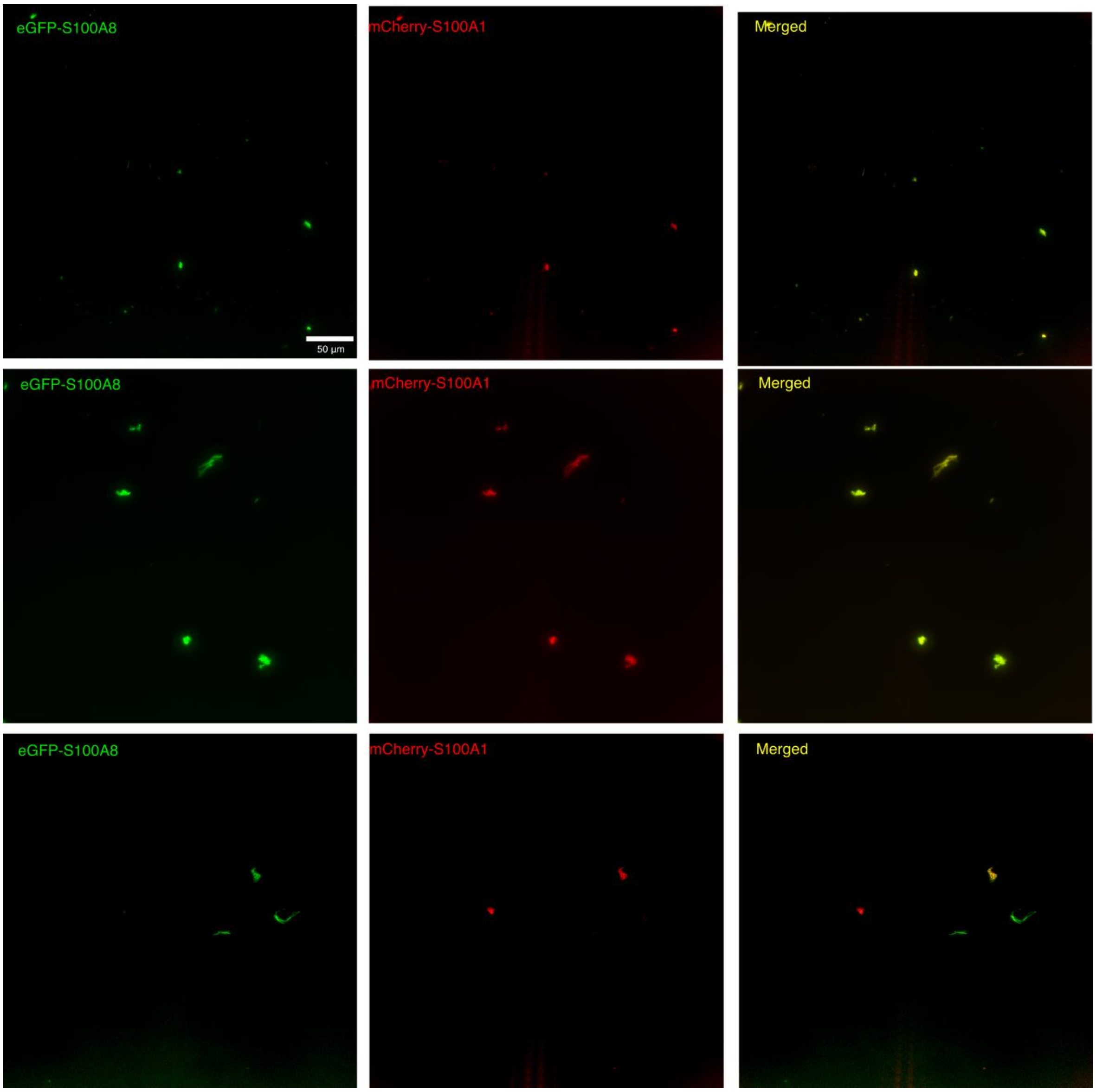


**Figure S4.** Fluorescence microscopy images of S100A1/S100A8 sample after aggregation in the presence of 1600 μM $CaCl_2$. Scale bar is 50 μm.

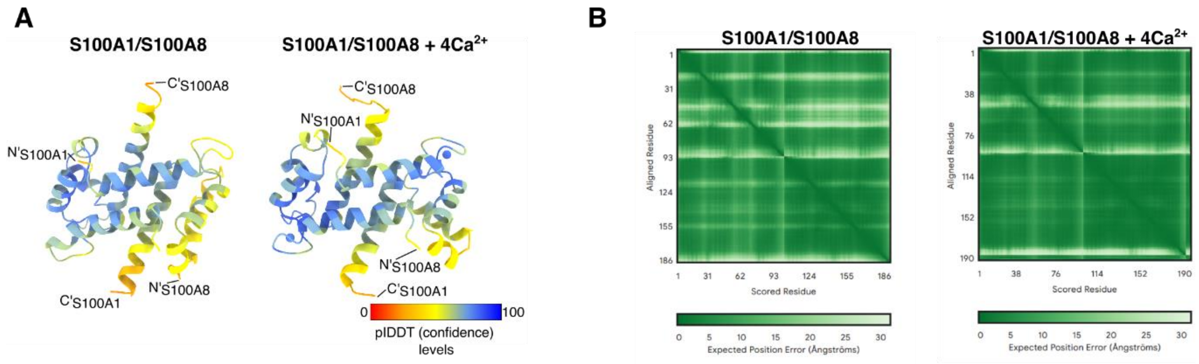


**Figure S5.** (**A**) Confidence levels of S100A1/S100A8 heterodimer with and without calcium ions. (**B**) 2D plots of Predicted Aligned Error, the darker green colour indicates lower position error of residue in structure.

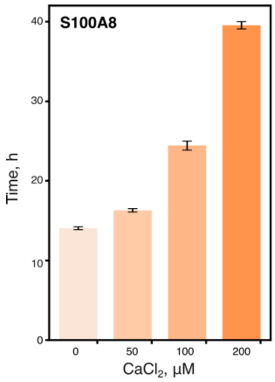


**Figure S6.** Inflection time of the first phase of S100A8 aggregation curves at different $CaCl_2$ concentrations.

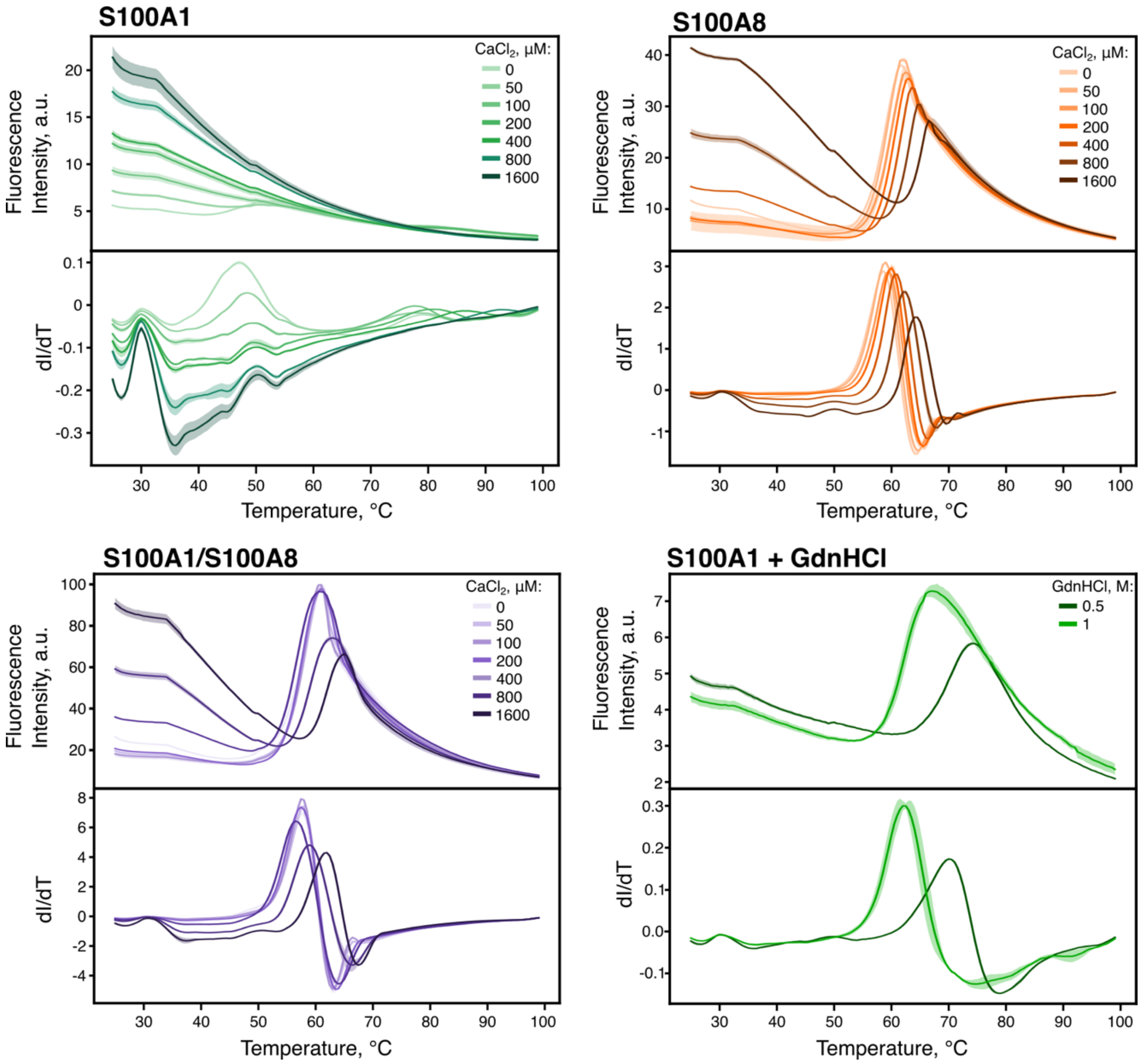


**Figure S7.** Thermal denaturation profiles and their first derivatives of S100A1, S100A8 and S100A1/100A8 proteins.

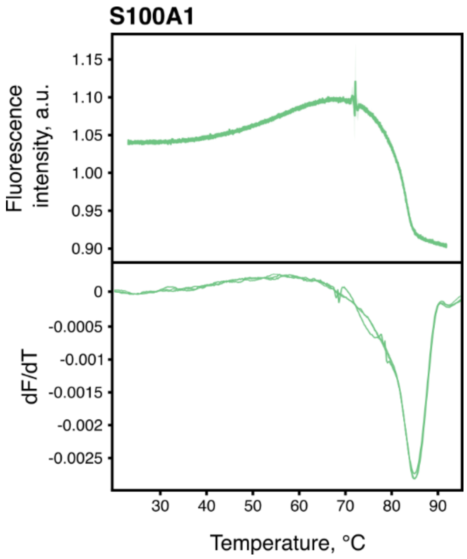


**Figure S8.** Thermal denaturation profiles and their first derivative of S100A1 using nanoDSF.

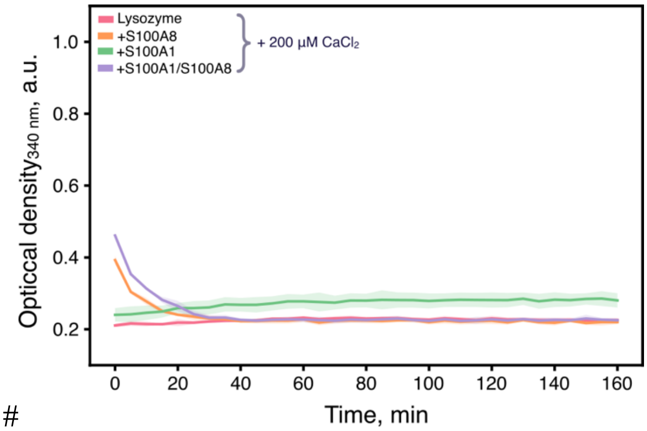


#

**Figure S9.** Chaperone activity assay of S100A1 and S100A8 (10 µM each) against DTT-induced aggregation of lysozyme (0.2 mg/ml) in the presence of 200 µM $CaCl_2$.

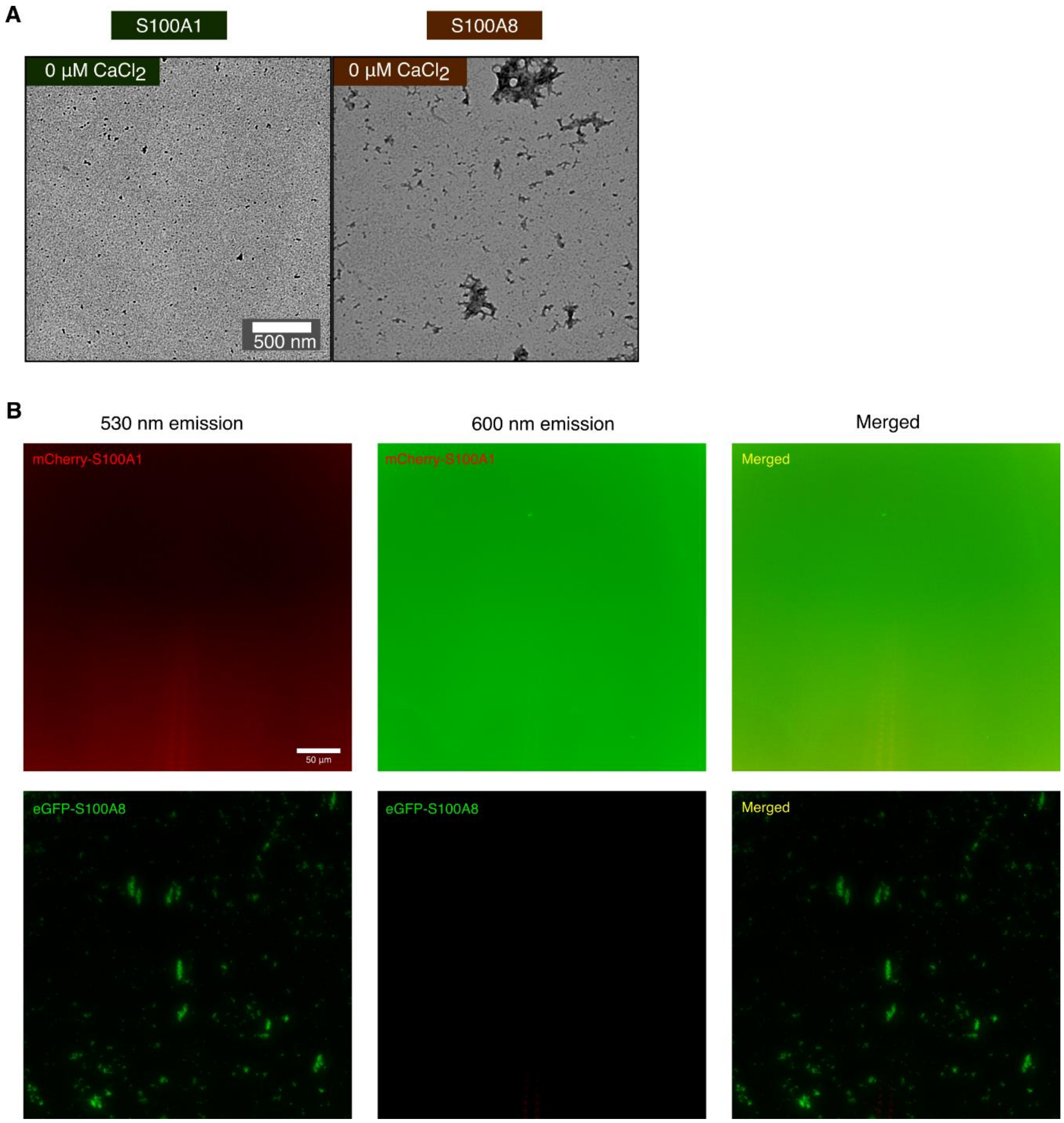


**Figure S10. (A)** TEM and (**B**) fluorescence microscopy images of S100A1 and S1008 samples after aggregation. Scale bars are 500 nm and 5 µm, respectively. Fluorescence levels are adjusted to the same level for each protein and emission.

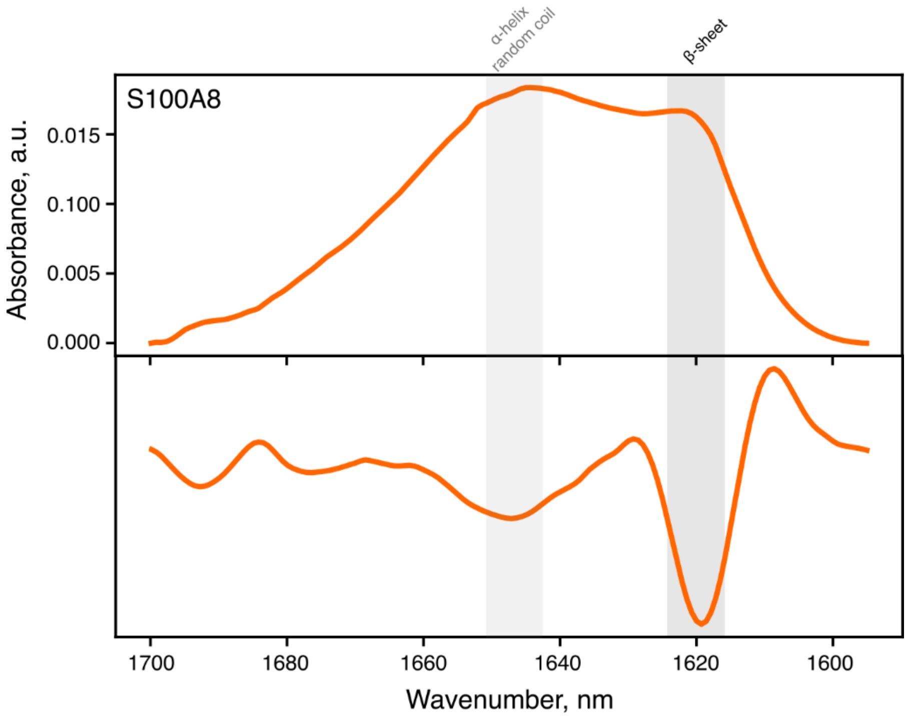


**Figure S11.** FTIR spectrum and its second derivative of S100A8 protein after 65 h of aggregation at 42 °C.

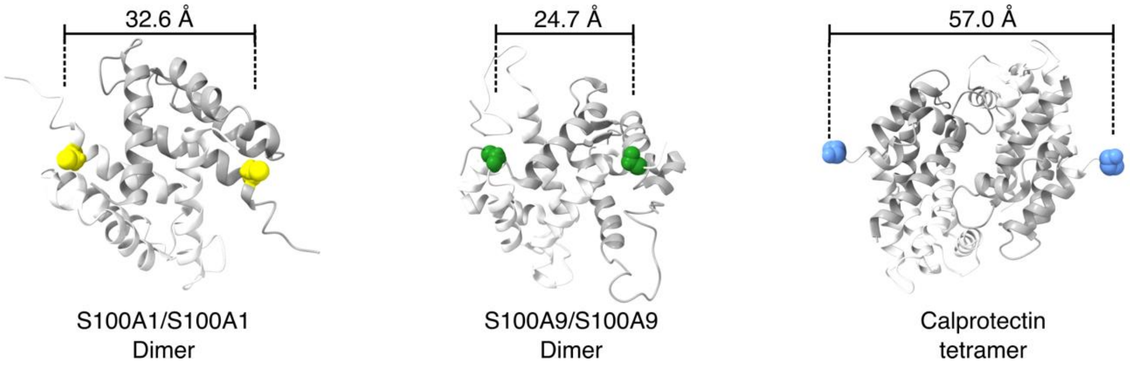


**Figure S12.** Measured distances between colored (spheres) residues in S100A1 dimer (PDB id: 2l0P), S100A9 dimer (PDB id: 5I8N) and tetramer structure of calprotectin (PDB id:1XK4). The colour residues correspond to the closest available positions to labelled cysteine with MTSSL.

| Plasmid | Primer | Sequence |
|---|---|---|
| pVK1<br>(eGFP-S100A8) | GFP_S100A8_rf_frw<br>GFP_S100A8_rf_rev | 5' TGGGCCATCACCATCACCATCACGTGAGCAAGGGCGAGGAG 3'<br>5' CGCTTTCTCCAGCTCGGTCAACATCGAAGCTTGAGCTCGAGA 3' |
| pVK2<br>(mCherry-S100A1) | mCherry_S100A1_rf_frw<br>mCherry_S100A1_rf_rev | 5' CCTGGTGCCGCGCGGCAGCCATATGGTGAGCAAGGGCGAA 3'<br>5' TCCATAGCTGTCTCTAGTTCTGATCCCATAGAACCACCACCACCAGA 3' |

Supplementary Table 1: Plasmids and respective primers used in this study.
